# The substrate scopes of enzymes: a general prediction model based on machine and deep learning

**DOI:** 10.1101/2022.05.24.493213

**Authors:** Alexander Kroll, Sahasra Ranjan, Martin K. M. Engqvist, Martin J. Lercher

## Abstract

For a comprehensive understanding of metabolism, it is necessary to know all potential substrates for each enzyme encoded in an organism’s genome. However, for most proteins annotated as enzymes, it is unknown which primary and/or secondary reactions they catalyze [1], as experimental characterizations are time-consuming and costly. Machine learning predictions could provide an efficient alternative, but are hampered by a lack of information regarding enzyme non-substrates, as available training data comprises mainly positive examples. Here, we present ESP, a general machine learning model for the prediction of enzyme-substrate pairs, with an accuracy of over 90% on independent and diverse test data. This accuracy was achieved by representing enzymes through a modified transformer model [2] with a trained, task-specific token, and by augmenting the positive training data by randomly sampling small molecules and assigning them as non-substrates. ESP can be applied successfully across widely different enzymes and a broad range of metabolites. It outperforms recently published models designed for individual, well-studied enzyme families, which use much more detailed input data [3, 4]. We implemented a user-friendly web server to predict the substrate scope of arbitrary enzymes, which may support not only basic science, but also the development of pharmaceuticals and bioengineering processes.

## Introduction

Enzymes not only evolved to efficiently catalyze one or more specific chemical reactions, increasing their reaction rates up to over a million-fold over the spontaneous rates^5^. In addition, most enzymes are promiscuous, i.e., they catalyze further, physiologically irrelevant or even harmful reactions^6,7,8^. Accordingly, a comprehensive mapping of enzyme-substrate relationships plays a crucial role in pharmaceutical research and bio-engineering, e.g., for the production of drugs, chemicals, food, and biofuels^9,10,11^.

Unfortunately, it is both expensive and time-consuming to determine experimentally which reactions are catalyzed by a given enzyme. There is thus a huge imbalance between the number of proteins predicted to be enzymes and the experimental knowledge about their substrate scopes. While the UniProt database^1^ contains entries for over 36 million different enzymes, more than 99% of these lack high quality annotations of the catalyzed reactions. Efforts are underway to develop high-throughput methods for the experimental determination of enzyme-substrate relationships, but these are still in their infancy^12,13,14^.

Furthermore, even high-throughput methods cannot deal with the vast search space of all possible small molecule substrates, but require the experimenter to choose a small subset for testing.

Our goal in this study was to develop a single machine learning model capable of predicting enzyme-substrate relationships across all proteins, thereby providing a tool that helps to focus experimental efforts on enzyme-small molecule pairs likely to be biologically relevant. Developing such a model faces two major challenges. First, a numerical representation of each enzyme that is maximally informative for the downstream prediction task must be obtained^15^. To be as broadly applicable as possible, these representations should be based solely on the enzymes’ primary sequence and not require additional features, such as binding site characteristics. Second, public enzyme databases only list positive instances, i.e., molecules with which enzymes display measurable activity (substrates)^16^. For training a prediction model, an automated strategy for obtaining negative, non-binding enzyme-small molecule instances must thus be devised.

Existing machine learning approaches for predicting enzyme-substrate pairs were developed specifically for small enzyme families for which unusually comprehensive training datasets are available^3,4,16,17,18^. For example, Mou et al.^3^ developed models to predict the substrates of bacterial nitrilases, using input features based on the 3D-structures and active sites of the enzymes. They trained various machine learning models based on experimental evidence for all possible enzyme-small molecule combinations within the models’ prediction scope (*N* = 240). Yang et al.^4^ followed a similar approach, predicting the substrate scope of plant glycosyltransferases among a pre-defined set of small molecules. They trained a decision tree-based model with a dataset covering almost all possible enzyme-small molecule combinations. Pertusi et al.^16^ trained four different support vectors machines (SVMs), each for a specific enzyme. As input features, their models only use information about the (potential) substrates, as well as non-substrates manually extracted from the literature; no explicit information about the enzymes was used. Roettig et al.^17^ and Chevrette et al.^18^ predicted the substrate scopes of small enzyme families, training machine learning models with structural information relating to the enzymes active sites.

All these models aim to predict substrates for one single enzyme or for a small group of enzymes. In each case, the approach relies on very dense experimental training data, i.e., experimental results for all or almost all potential enzyme-substrate pairs. However, for the vast majority of enzyme families, such extensive training data is not available. As yet, there have been no published attempts to formulate and train a general model that can be applied across widely different enzyme families.

Deep learning models have been used to predict enzyme functions by either predicting their assignment to EC classes^19,20,21^, or by predicting functional domains within the protein sequence^22^. These approaches are complementary to the prediction of enzyme-substrate pairs, as the substrate scopes of different enzymes within a given EC class or with a specific domain architecture can be highly diverse^23^.

Prior work related to the prediction of enzyme-substrate pairs are the prediction of drug-target binding affinities (DTBAs) and of Michaelis-Menten constants, *K*_M_ and *k*_cat_. State-of-the-art approaches in this domain are feature-based, i.e., numerical representations of the protein and the substrate molecule are used as input to machine learning models^24,25,26,27,28^. As numerical representations for the substrate molecule, these approaches use SMILES representations^29^, expert-crafted fingerprints^30^, or fingerprints created with graph neural networks^31,32^. Proteins are usually encoded numerically through deep learning-based representations of the amino acid sequences^2,33,34^. However, these approaches cannot be transferred one-to-one to the problem of predicting enzyme-substrate pairs. The *K*_M_ and *k*_cat_ prediction models are exclusively trained with positive enzyme-substrate pairs and therefore cannot classify molecules as substrates or non-substrates^27,28^. Many of the proteins used to train the DTBA prediction models have no enzymatic functions; even if they do, the molecules used for training are mostly not naturally occurring potential substrates, and thus there has been no natural selection for or against binding. In contrast, the binding between enzymes and substrates evolved under natural selection. It appears likely that this evolutionary relationship influences our ability to predict enzyme-substrate pairs, and DTBA models are thus not expected to perform well at this task.

Here, we go beyond the current state-of-the-art by creating maximally informative protein representations, using a customized, task-specific version of the *ESM-1b* transformer model^2^. The model contains an extra 1280-dimensional token, which was trained end-to-end to store enzyme-related information salient to the downstream prediction task. This general approach was first introduced for natural language processing^35^, but has not yet been applied to protein feature prediction. We created negative training examples using data augmentation, by randomly sampling small molecules structurally similar to the substrates in experimentally confirmed enzyme-substrate pairs. This approach was guided by the biochemical intuition that enzymes are specific catalysts; for any given enzyme, the vast majority of randomly selected small molecules will not be substrates. Small molecules were represented numerically using extended-connectivity fingerprints (ECFPs)^30^. A gradient-boosted decision tree model was trained on the protein and small molecule representations for a high-quality dataset with ∼ 18000 very diverse, experimentally confirmed positive enzyme-substrate pairs (**Figure 1**). The resulting Enzyme Substrate Prediction model – ESP – achieves high prediction accuracy and outperforms previously published enzyme family-specific prediction models. This work demonstrates how trained task-specific tokens and augmented datasets can be used to overcome challenges in predicting enzyme-substrate relationships.

**Figure 1.**
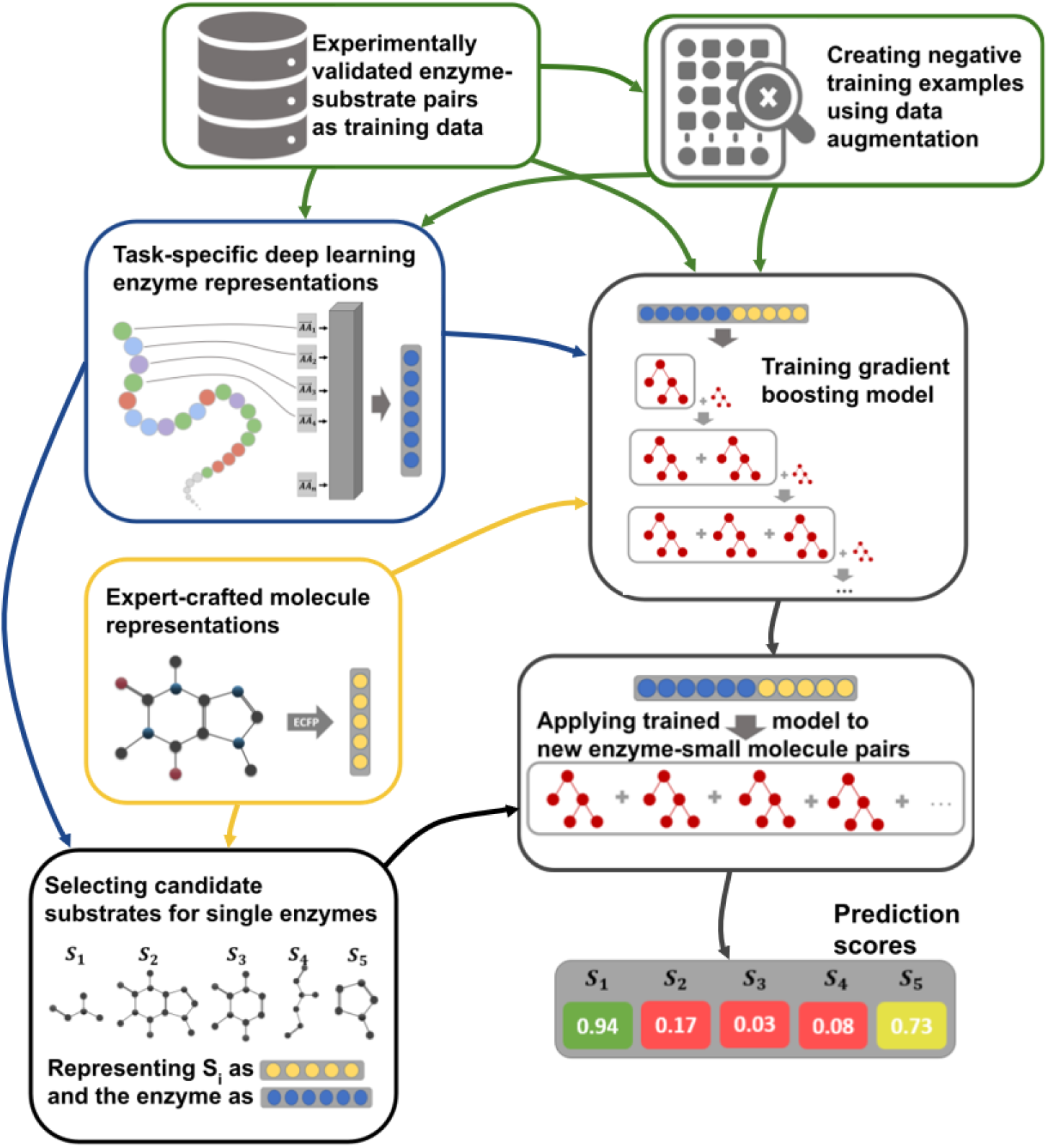
Model overview. Experimentally validated enzyme-substrate pairs and sampled negative enzyme-small metabolite pairs are numerically represented with task-specific enzyme representations and expert-crafted small metabolite representations. Concatenated enzyme-small metabolite representations are used to train a gradient boosting model. After training, the fitted model can be used to predict promising candidate substrates for enzymes.

## Results

### Obtaining training and test data

We created a dataset with experimentally confirmed enzyme-substrate pairs using the GO annotation database for UniProt IDs^36^ (Methods, “Creating a database with enzyme-substrate pairs”). We were able to extract 18351 enzyme-substrate pairs with experimental evidence for binding, containing 12156 unique enzymes and 1379 unique metabolites. We also extracted 274030 enzyme-substrate pairs with phylogenetically inferred evidence, i.e., these enzymes are closely related to enzymes associated with the same reactions. These ”guilt by association” assignments are much less reliable than direct experimental evidence. Thus, we only used these additional data during pre-training to create task-specific enzyme representations, but did not use them for final model training.

There is no systematic information on negative enzyme-small molecule pairs, i.e., pairs where the molecule is not a substrate of the enzyme. We hypothesized that such negative data points could be created artificially through random sampling. Enzymes are specific catalysts, and a randomly sampled small molecule therefore appears unlikely to be a substrate. For a small, but ultimately unknown, number of enzyme-small molecule pairs this assumption will result in incorrect negative labels. *A priori*, the frequency of this occurrence was deemed sufficiently low to not adversely affect model performance. This assumption was confirmed *a posteriori* by the high model accuracy on independent test data (see below).

To force the model to learn to distinguish between similar potential substrates, we aimed to sample small metabolites as negative data points that are structurally similar to the true substrates – so-called hard negatives. We used a similarity score based on molecular fingerprints, with values ranging from 0 (no similarity) to 1 (identity; see Methods, “Sampling negative data points”). For every positive enzyme-substrate pair, we sampled three molecules with similarity scores between 0.75 and 0.95 to the actual substrate of the enzyme, and used them to construct negative enzyme-molecule pairs. We opted for creating more negative data points than we have positive data points, as this not only provided us with more data, but it also more closely reflects the true distribution of positive and negative data points compared to a balanced distribution.

Our final dataset comprises 69365 entries. We split this data into a training set (80%) and a test set (20%). We made sure that no enzyme in the test set has a sequence identity higher than 80% compared to any enzyme in the training set, as the prediction task would be too easy if two very similar enzyme-small metabolite pairs existed in the training and test sets.

### Representing small molecules as extended-connectivity fingerprints

Extended-connectivity fingerprints (ECFPs) are binary representations for small molecules, which are represented as graphs; atoms are interpreted as nodes and chemical bonds as edges. Bond types and feature vectors with information about every atom are calculated (types, masses, valences, atomic numbers, atom charges, and number of attached hydrogen atoms)^30^. Afterwards, these identifiers are updated for a fixed number of steps by iteratively applying predefined functions to summarize aspects of neighboring atoms and bonds. After the iteration process, all identifiers are converted into a single binary vector with structural information about the molecule. The number of iterations and the dimension of the fingerprint can be chosen freely. We set them to the default values of 3 and 1024, respectively; lower or higher dimensions led to inferior predictions.

### Representing enzymes through a modified state-of-the-art deep learning architecture

The *ESM-1b* model is a state-of-the-art transformer network^37^, trained with ∼27 million proteins from the UniRef50 dataset^38^ in a self-supervised fashion^2^. This model takes an amino acid sequence as its input and outputs a numerical representation of the sequence; these representations are often referred to as protein embeddings. During training of *ESM-1b*, ∼ 15% of the amino acids in a protein’s sequence are randomly masked and the model is trained to predict the identity of the masked amino acids (see **Figure 2a**). This training procedure forces the model to store both local and global information about the protein sequence in one 1280-dimensional representation vector for each individual amino acid. In order to create a single fixed-length numerical representation of the whole protein, one typically calculates the element-wise mean across all amino acid representations^33,2,39^. We refer to these as *ESM-1b* vectors.

**Figure 2.**
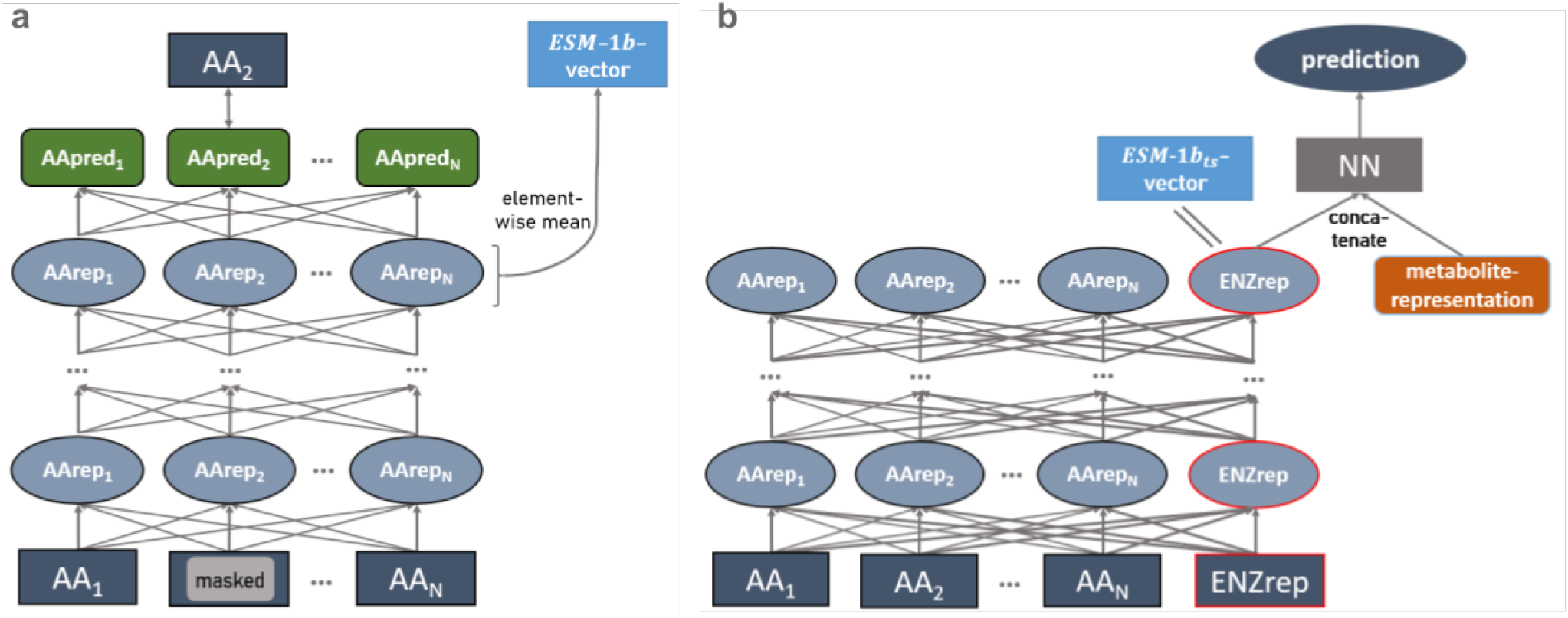
A task-specific enzyme representation developed from the *ESM-1b* model. **(a)** *ESM-1b* model. Amino acids of a protein sequence are represented with numerical vectors and passed through a transformer network. Some amino acid representations are masked. All representations are iteratively updated 33 times, using information about neighboring and distant amino acids. The *ESM-1b* model is trained to predict the masked amino acids. *ESM-1b* vectors are calculated by taking the element-wise mean of all representations in the last layer. **(b)** Modified *ESM-1b* model. An additional representation for the whole enzyme is added to the amino acid representations. After updating all representations 33 times, the enzyme representation is concatenated with a small molecule representation. The network is trained to predict whether the small molecule is a substrate for the given enzyme. After training, the *ESM-1b*_*ts*_ vector is extracted as the enzyme representation before adding the small molecule representation.

However, simply taking the element-wise mean results in information loss and does not consider the task for which the representations shall be used, which can lead to subpar performance^15^. To overcome these issues, we created task-specific enzyme representations optimized for the prediction of enzyme-substrate pairs. We slightly modified the architecture of the *ESM-1b* model, adding one additional 1280-dimensional enzyme representation in which to capture information salient to the downstream prediction task (**Figure 2b**). This enzyme representation was updated in the same way as the regular *ESM-1b* amino acid representations.

After a predefined number of update steps, the enzyme representation was concatenated with the small molecule ECFP-vector. The combined vector was used as the input for a fully connected neural network (FCNN), which was then trained end-to-end to predict whether the small molecule is a substrate for the enzyme. This approach facilitates the training of a single, optimized, task-specific representation. The *ESM-1b* model contains many parameters and thus requires substantial training data. Therefore, in the pre-training that produces the task-specific enzyme representations, we added phylogenetically inferred evidence to our training set; this resulated in ∼ 287000 data points. After pre-training, we used the network to extract the 1280-dimensional task-specific representations for all enzymes in our dataset. In the following, these representations are called *ESM-1b*_*ts*_ vectors.

### Successful prediction of enzyme-substrate pairs by using combined enzyme and small molecule representations

To compare the *ESM-1b* and *ESM-1b*_*ts*_ vectors, we investigated the performance of our machine learning models when combining either type with the small molecule representations. We concatenated one of the two 1280-dimensional enzyme representations and the 1024-dimensional ECFP vector, resulting in input vectors of dimension 2304. We used these inputs to train gradient boosted decision tree models^40^ for the binary classification task of predicting whether the metabolite is a substrate for the enzyme.

We performed hyperparameter optimizations for both models (trained with either *ESM-1b* or *ESM-1b*_*ts*_ vectors), including the parameters learning rate, depth of trees, number of iterations, and regularization coefficients. For this, we performed a random grid search with a 5-fold cross-validation (CV) on the training set (Methods, “Hyperparameter optimization of the gradient boosting models”). To account for the higher number of negative compared to positive training data, we also included a weight parameter that lowered the influence of the negative data points. The results of the cross-validations are displayed as boxplots in **Figure 3a**. The best sets of hyperparameters are displayed in **Suppl. Table S1**. After hyperparameter optimization, the models were trained with the best set of hyperparameters on the whole training set and were validated on our independent test set, which had not been used for model training or hyperparameter selection.

**Figure 3.**
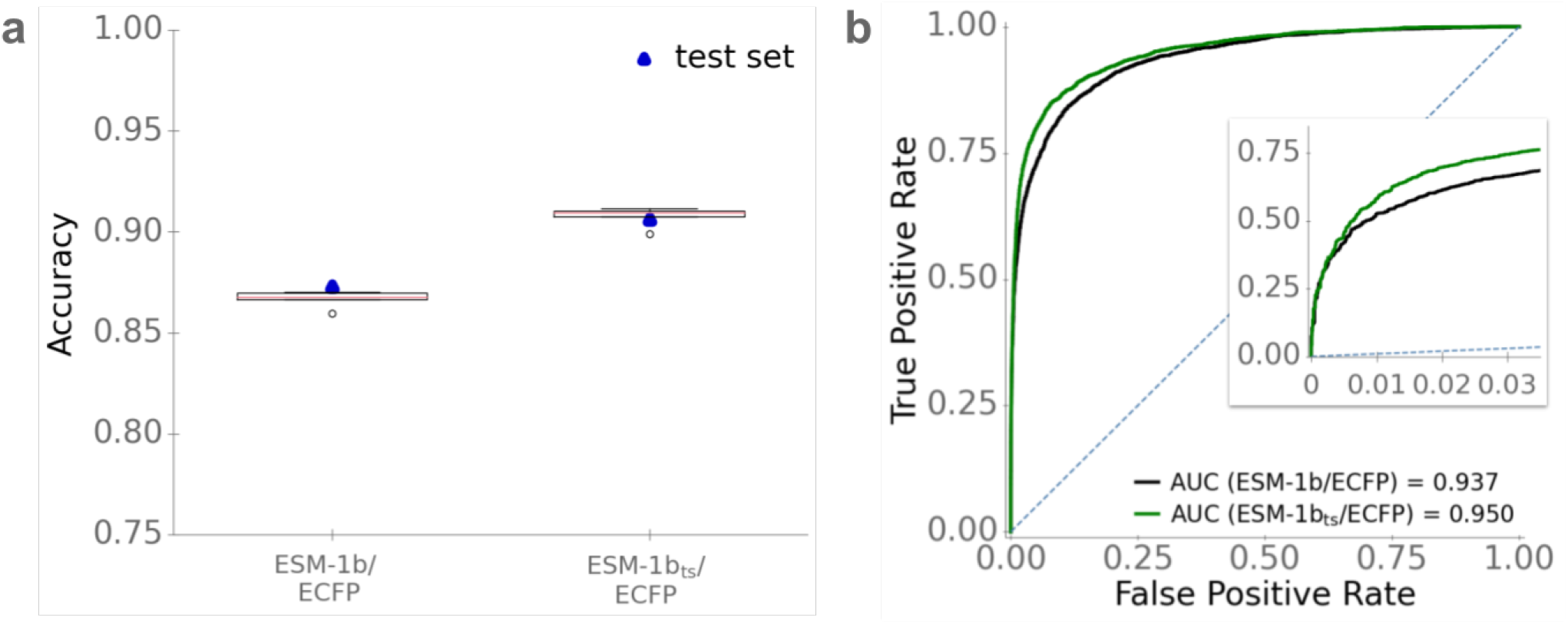
Optimized models provide accurate predictions of enzyme-substrate pairs. **(a)** Accuracies. Boxplots summarize the results of the 5-fold CV on the training set with the best sets of hyperparameters. Blue dots display the accuracies on the test set, using the optimized models trained on the whole training set. **(b)** ROC curves for the test set. The dotted line displays the ROC curve expected for a completely random model.

Commonly used metrics to measure the performance of binary classification models are accuracy, ROC-AUC score, and Matthews correlation coefficient (MCC). Accuracy is simply the fraction of correctly predicted data points among the test data. The ROC-AUC score is a value between 0 and 1 that summarizes how well a classifier is able to distinguish between the positive and negative class, where a value of 0.5 would result from a model that randomly assigns class labels, and a value of 1 corresponds to perfect predictions. The MCC is a correlation coefficient for binary data, comparable to the Pearson correlation coefficient for continuous data; it takes values between -1 and +1, where 0 would result from a model that randomly assigns class labels, and +1 indicates perfect agreement.

The model that we trained with *ESM-1b* vectors achieves an accuracy of 87.5%, a ROC-AUC score of 0.937, and an MCC of 0.70. The model trained with *ESM-1b*_*ts*_ vectors was substantially better, achieving an accuracy of 90.6%, a ROC-AUC score of 0.950, and an MCC of 0.76 (**Figure 3**); this difference is statistically highly significant (McNemar’s test: *P <* 10^−28^).

We also trained gradient boosting models with task-specific small molecule representations that we created with graph neural networks (GNNs; see Methods, “Calculating task-specific fingerprints for the metabolites using graph neural networks”). However, these led to very similar results (**Suppl. Table S1**) and are far more laborious to calculate. As the best Enzyme Substrate Prediction model (ESP), we thus chose the model based on *ESM-1b*_*ts*_ for enzymes and ECFPs for small molecules.

### Good predictions even for enzymes with low sequence identity to training data

It appears likely that prediction quality is best when highly similar enzymes were in the training set, but decreases when no similar enzymes were used for training. How strong is that dependence? To answer this question, we first calculated the maximal enzyme sequence identity compared to the enzymes in the training set for all of the 2291 enzymes in the test set. Next, we split the test set into three subgroups: data points with enzymes with a maximal sequence identity between 0 and 40%, between 40% and 60%, and between 60% and 80%.

For data points with high sequence identity levels (60-80%), the ESP model is highly accurate, with accuracy 94%, ROC-AUC score 0.98, and MCC 0.85 (**Figure 4**). ESP still performs very well for data points with intermediate sequence identity levels (40-60%), achieving an accuracy of 92%, ROC-AUC-score 0.97, and MCC 0.81. Even for enzymes with low sequence identity to training data (0 − 40%), the ESP model achieves good results and classifies 88% of the data points correctly, with ROC-AUC score 0.92 and MCC 0.68. Thus, while using more similar enzymes during training improves the prediction quality, very good prediction accuracy can still be achieved for enzymes that are only distantly related to those in the training set. The observed differences were statistically significant for sequence identities 0-40% versus 40-60% (Mann–Whitney *U* test: *P <* 10^−10^), but not for 0-60% versus 60-80% (*P* = 0.137).

**Figure 4.**
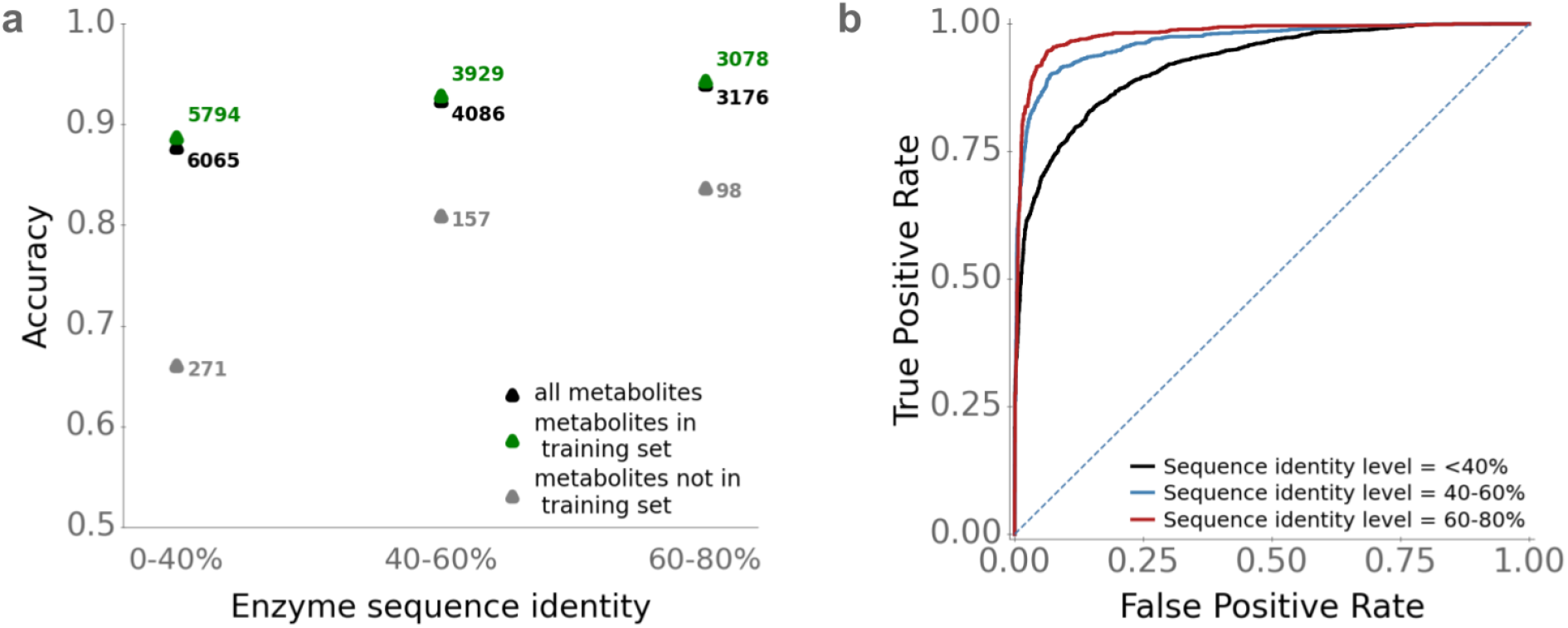
Accurate predictions for enzymes with 40-60% sequence identity to training data. We divided the test set into subsets with different levels of enzyme sequence identity compared to enzymes in the training set. **(a)** ESP accuracies, calculated separately for enzyme-small molecule pairs where the small molecule occurred in the training set and where it did not occur in the training set. **(b)** ESP ROC curves. The dotted line displays the ROC curve expected for a completely random model.

### Low model performance for unseen small molecules

In the previous subsection, we showed that model performance is highest for enzymes that are similar to proteins in the training set. Similarly, it appears likely that the model performs better when making predictions for small molecules that are also in the training set. To test this hypothesis, we divided the test set into data points with small molecules that occurred in the training set and those with small molecules that did not occur in the training set.

The ESP model does not perform well for data points with small molecules not present in the training set. When considering only enzyme-small molecules pairs with small molecules not represented in the training set and an enzyme sequence identity level of 0-40% compared to the training data, ESP achieves an accuracy of 66%, ROC-AUC score 0.57, and MCC 0.01. At an enzyme sequence identity level of 40-60%, accuracy improves to 80%, with ROC-AUC score 0.60, and MCC 0.02 for unseen small molecules. At high enzyme sequence identity levels of 60-80%, the accuracy reaches 84%, with ROC-AUC score 0.66, and MCC 0.12, still barely better than a model that makes completely random predictions. Thus, for unseen small molecules, even a very moderate model performance requires that proteins similar to the enzyme are present in the training set. We again found the differences to be statistically significant for 0-40% versus 40-60% (Mann–Whitney *U* test: *P <* 10^−7^), but not for 40-60% versus 60-80% (*P* = 0.167).

For those test data points with small molecules not present in the training set, we wondered if a high similarity of the small molecule compared to at least one substrate in the training set leads to improved predictions, analogous to what we observed for enzymes with higher sequence identities. For each small molecules not present in the training set, we calculated the maximal pairwise similarity score compared to all substrates in the training set. We could not find any evidence that a higher maximal similarity score leads to better model performance (**Suppl. Figure S1**). Hence, to achieve predictions with high accuracies for new enzyme-small molecule pairs, data points with the same small molecule should be present in the training set.

How many training data points with identical substrates are needed to achieve good model performance? For every small molecule in the test set, we counted how many times the same molecule is experimentally confirmed substrate in the training set. **Suppl. Figure S2** shows that having as few as two positive training data points for a given small molecule leads to good accuracy when pairing the same small molecule with other enzymes.

### Model performance increases with increased training set size

The previous subsections indicate that a bigger training set with a more diverse set of enzymes and small molecules should lead to improved performance. To test this hypothesis, we trained the gradient boosting model with different training set sizes, ranging from 30% to 100% of the available training data. **Figure 5** shows that accuracy and ROC-AUC score indeed increase with increasing training set size (Spearman rank correlations, accuracy: *ρ*^2^ = 0.95, *P <* 10^−4^; ROC-AUC score: *ρ*^2^ = 1.0, *P <* 10^−15^).

**Figure 5.**
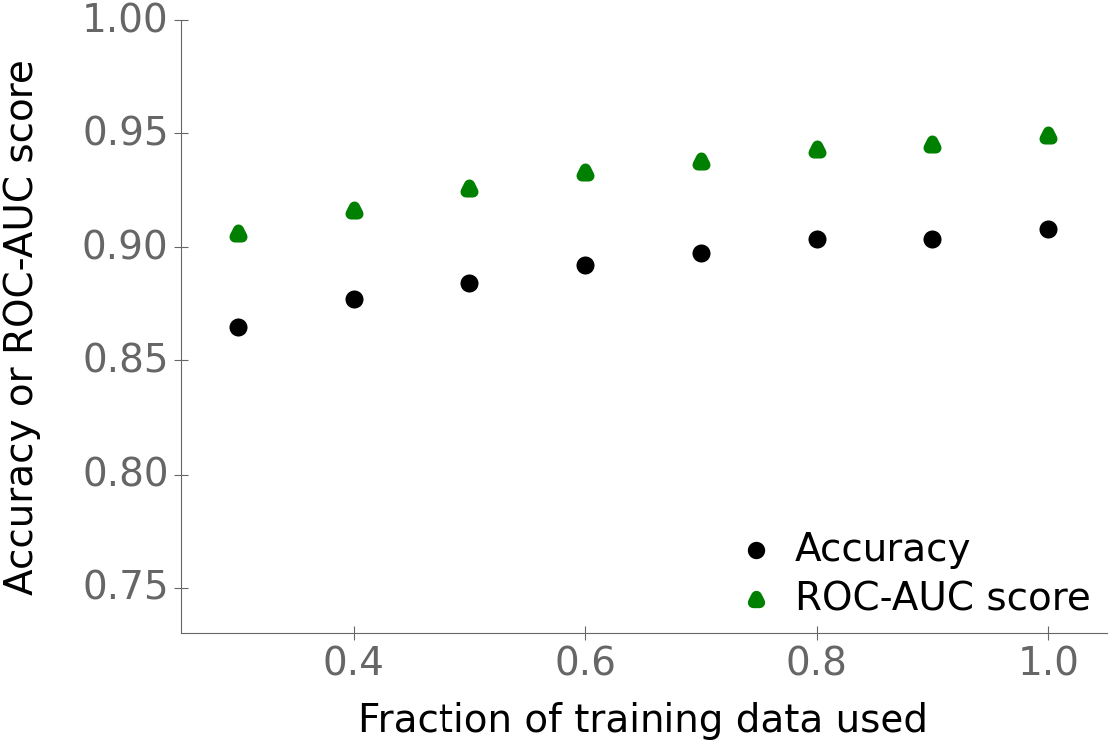
Model performance increases with training set size. Points show accuracies and ROC-AUC scores for the test set versus the fraction of the available training data used for training the gradient boosting model.

### ESP is able to say “I don’t know” for some data points

Our trained classification model does not simply output the positive or negative class as a prediction. Instead, it outputs a prediction score between 0 and 1, which can be interpreted as a measurement of the probability for a data point to belong to the positive class. So far, we assigned all predictions with a score ≥ 0.5 to the positive class, and all predictions below 0.5 to the negative class. **Figure 6** displays the distributions of the true (blue) and false (red) predictions for our test set across prediction scores.

**Figure 6.**
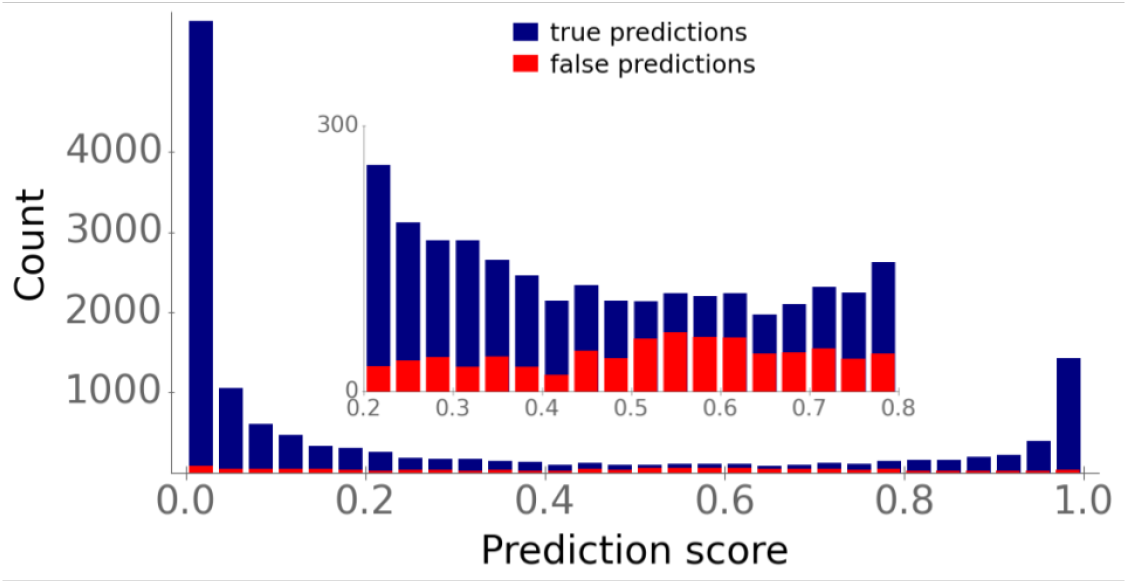
Prediction scores around 0.5 indicate model uncertainty. Stacked histogram bars display the prediction score distributions of true predictions (blue) and false predictions (red). The inset shows a blow-up of the interval [0.2, 0.8].

Most true predictions have a score either close to 0 or close to 1, i.e., the ESP model is very confident about these predictions. In contrast, false predictions are distributed much more evenly across prediction scores. Approximately 5% of prediction scores for our test data falls between 0.4 and 0.6. The model seems to be uncertain for these data points: for this subset, predictions are only barely better than random predictions, with an accuracy of 55%, ROC-AUC score 0.58, and MCC 0.10. Thus, when applied in practice, prediction scores between 0.4 and 0.6 should be considered uncertain and should not be assigned to one of the two classes.

### ESP outperforms two recently published models for predicting the substrate scope of enzymes

We compared ESP with two recently published models for predicting the substrate scope of specific enzyme families. Mou et al.^3^ published a machine learning model for predicting substrates of bacterial nitrilases. For model training and validation, they used a dataset with all possible combinations of 12 enzymes and 20 small molecules (*N* = 240), randomly split into 80% training data and 20% test data. We added all training data from Ref.^3^ to our training set and validated the updated ESP model on the corresponding test data, which had no overlap with our training data. Mou et al.^3^ achieved an accuracy of 82% and a ROC-AUC score of 0.90 on the test set. ESP achieves better results, with an accuracy of 87.5%, ROC-AUC score 0.94, and MCC 0.75. This improvement is particularly striking given that Mou et al.^3^ used knowledge about the enzymes’ 3D structures and binding sites, while we only use a representation of the linear amino acid sequence.

Yang et al.^4^ published a decision tree-based model, GT-Predict, for predicting the substrate scope of glycosyltransferases of plants. As a training set, they used 2847 data points with 59 different small molecules and 53 different enzymes from *Arabidopsis thaliana*, i.e., the data covered 90.7% of all possible enzyme-small molecule combinations. These authors used two independent test sets to validate the model, one dataset with 266 data points with enzymes from *Avena strigosa* and another dataset with 380 data points with enzymes from *Lycium barbarum*. On those two test sets, GT-Predict achieves accuracies of 79.0% and 78.8%, respectively, and MCCs of 0.338 and 0.319, respectively. We added the training set from Ref.^4^ to our training set. The test sets from *Avena strigosa* and *Lycium barbarum* had no overlap with our training data. For these two sets, we achieved similar accuracies of 78.2% and 78.2%, respectively, ROC-AUC scores of 0.80 and 0.84, respectively, and improved MCCs of 0.484 and 0.517, respectively. As the datasets by Yang et al.^4^ are imbalanced with a proportion of 18-31% of positive data points, the MCC is a more meaningful score compared to the accuracy^41^, and hence we conclude that ESP outperforms GT-Predict.

Comparison of our model predictions to these two (almost) complete experimental datasets indicates that ESP is indeed capable of predicting the full substrate scope of enzymes.

We also tested model performances for the test sets by Mou et al.^3^ and Yang et al.^4^ without adding any new training data to the ESP. Only ∼ 5% and ∼ 8% of the small molecules in these test sets did already occur in our training set. Since we have shown above that model performance massively drops if the model is applied to unseen small molecules (**Figure 4a**), we did not expect good model performances. Indeed, for all three test sets, accuracies are below 68%, ROC-AUC scores are below 0.59, and MCCs are below 0.12 (**Suppl. Table S2**).

### The ESP web server facilitates an easy use of the prediction model

We implemented a web server that allows an easy use of ESP without requiring programming skills or the installation of specialized software. It is available at https://esp.cs.hhu.de. As input, the web server requires an enzyme amino acid sequence and a representation of a small molecule (either as a SMILES string, KEGG Compound ID, or InChI string). Users can either enter a single enzyme-small molecule pair into an online form, or upload a CSV file with multiple pairs. In addition to the prediction score, the ESP web server reports how often the entered metabolite was present as a true substrate in our training set. Since we have shown that model performance massively drops when the model is applied to new small molecules, we recommend to use the prediction tool only for those small molecules represented in our training set. We uploaded a full list with all small molecules from the training set to the web server homepage, listing how often each one is present among the positive data points.

## Discussion

We presented a general approach for predicting the substrate scope of enzymes; ESP achieves an accuracy of over 90% on an independent test set with enzymes that share at most 80% sequence identity with any enzyme used for training. Notably, the model performs with an accuracy of 88% even for enzymes with very low sequence identity (*<* 40%) to proteins in the training set. This performance seems remarkable, as it is believed that enzymes often evolve different substrate specificities or even different functions if sequence identity falls below 40%^42^.

To achieve these results, we used very general input features: an expert-crafted fingerprint with structural information about the small molecule^30^ and a numerical representation of the enzyme calculated from its amino acid sequence^2^. We showed that creating task-specific enzyme representations leads to significant improvements compared to non-task-specific enzyme representations (**Figure 3**). Moreover, our results clearly show that randomly sampling negative enzyme-molecule pairs is an effective and viable approach. Future refinements of this approach might boost model performance further.

Despite the structural similarities of ESP to state-of-the-art models for predicting drug–target binding affinities (DTBAs) and for predicting Michaelis-Menten constants of enzyme-substrate pairs^24,25,26,27,28^, the performances of these models are not comparable, as we trained ESP for a binary classification task, whereas the other models address regression tasks. Instead, we compared our approach to two recently published models for predicting enzyme-substrate pairs^3,4^. These two models used very specific input features, such as an enzyme’s active site properties and physicochemical properties of the metabolite, and were designed and trained for only a single enzyme family. Our general ESP model achieves superior results, despite learning and extracting all relevant information for this task from much less detailed, general input representations. The application of ESP to the dataset from Mou et al.^3^ also demonstrated that our model can successfully distinguish between similar potential substrates for the same enzyme, as it achieved good results when it was applied to different nitriles for bacterial nitrilases.

One limitation of ESP is that model performance massively drops for small molecules that did not occur in the training set. However, the current version of ESP can still be applied successfully to a broad range of almost 1400 different small molecules present in our dataset. Once more training data becomes available, model performance will very likely improve further (**Figure 5**). Mining other biochemical databases, such as BRENDA^43^, might be a low-cost avenue to expanding the number of different small molecules in the dataset. Adding as few as two additional positive training data points for new molecules will typically lead to very accurate predictions (**Suppl. Figure S2**).

The recent development of AlphaFold^44^ and RoseTTAFold^45^ facilitates predictions of the 3D structure for any protein with known amino acid sequence. Future work may also include input features extracted from such predicted enzyme structures. Our high-quality dataset with many positive and negative enzyme-small metabolite pairs, which is available on GitHub, might be a promising starting point to explore the utility of such features.

A main use case for the ESP model will be the prediction of possible substrate candidates for single enzymes. In contrast, ESP will likely not lead to satisfactory results when used to predict all enzyme-substrate pairs in a genome scale metabolic model. This problem results from the trade-off between the True Positive Rate (TPR) and the False Postive Rate (FPR) for different classification thresholds (**Figure 3b**). For example, choosing a classification threshold with a TPR of ∼ 80% leads to a FPR of ∼ 5%. If we consider a genome scale model with approximately 2000 enzymes and 2000 metabolites, then there exist ∼ 4 × 10^6^ possible enzyme-small molecule pairs, of which only about 6000 will be true enzyme-substrate pairs. A TPR of 80% would lead to the successful detection of 4800 true pairs. At the same time, an FPR of 5% would lead to an additional ∼ 200000 false predictions.

If, on the other hand, ESP is applied to a set of pre-selected candidate substrates for a single enzyme, a false positive rate of 5% can be acceptable. If we choose 200 molecules as substrate candidates, where one of these 200 is a true substrate for the enzyme, an FPR of 5 % means that the model predicts only ∼ 10 molecules falsely as a substrate, and there is an 80% chance that the true substrate is labeled correctly. This could help to bring down the experimental burden – and associated costs – of biochemical assays to levels where laboratory tests become tractable.

## Methods

### Software and code availability

All software was coded in Python^46^. We implemented and trained the neural networks using the deep learning library PyTorch^47^. We fitted the gradient boosting models using the library XGBoost^40^. The Python code used to generate the results, in the form of Jupyter notebooks, as well as all datasets, are available from https://github.com/AlexanderKroll/SubFinder.

### Creating a database with enzyme-substrate pairs

To create a database with positive enzyme-substrate pairs, we searched the Gene Ontology (GO) annotation database for UniProt IDs^36^ with over 922 million entries for annotations of the catalytic activity of enzymes. A GO annotation consists of a GO Term that is assigned to a UniProt ID, which is an identifier for proteins. GO Terms can contain information about the biological processes, molecular functions, and cellular components of proteins^48^. We first created a list with all catalytic GO Terms containing information about enzyme-catalyzed reactions. For each of these GO Terms, we extracted identifiers for the substrates involved in the reaction. For this purpose, we used a RHEA reaction ID^49^ from the GO Term, if available, or we extracted the substrate names via text mining from the definition of the GO Term. Substrate names were then mapped to KEGG and ChEBI identifiers via the synonym database from KEGG^50^, or, if no entry in KEGG was found, the PubChem synonym database^51^. We discarded all catalytic GO Terms for which we could not map at least one substrate to an identifier.

Entries in the GO annotation database have different levels of evidence: experimental, phylogenetically-inferred, computational analysis, author statement, curator statement, and electronic evidence. We searched the database for entries with catalytic GO Terms with either experimental or phylogenetically-inferred evidence. We extracted protein and substrate IDs from these entries. We removed all data points with water, oxygen, and ions, as these small substrates did not lead to unique ECFP representations (see below). We found 18351 enzyme-substrate pairs with experimental evidence. Among these data points, we have 12156 unique enzymes and 1379 unique substrates. We also found 274030 enzyme-substrate pairs with phylogenetically-inferred evidence. Among these data points are 198259 unique enzymes and 661 unique substrates. We downloaded the amino acid sequences for all enzymes via the UniProt mapping service^1^.

### Sampling negative data points

For every positive enzyme-substrate pair in our dataset, we created three negative data points for the same enzyme by sampling small molecules that are similar to the substrate of the positive data point. For this purpose, we first calculated the pairwise similarity of all small molecules in our dataset with the function FingerprintSimilarity from the RDKit package DataStructs^52^. This function uses molecular fingerprints of the molecules as its input and computes values between zero (no similarity) and one (high similarity). If possible, we sampled small molecules with a similarity score between 0.7 and 0.95. If we did not find such molecules, we reduced the lower bound in steps of 0.2 until enough small molecules could be sampled. During this sampling process, we took the distribution of the small molecules among the positive data points into account, i.e., molecules that occur more frequently as substrates among the positive data points also appear more frequently among the negative data points. To achieve this, we excluded small molecules from the sampling process if these molecules were already sampled enough times.

### Splitting the dataset into training and test sets

Before we split the dataset into training and test sets, we clustered all sequences by amino acid sequence identity using the CD-HIT algorithm^53^. We split the dataset randomly into 80% training data and 20% test data using a sequence identity cutoff of 80%, i.e., every enzyme in the test set has a maximal sequence identity of 80% compared to any enzyme in the training set. To analyze the ESP performance for different sequence identity levels, we further split the test set into subsets with sequence identity levels of 0-40%, 40-60%, and 60-80% using the CD-HIT algorithm^53^.

### Calculating extended-connectivity fingerprints for the small molecules

For every small molecule with a KEGG ID in our final dataset, we downloaded an MDL Molfile with 2D projections of its atoms and bonds from KEGG^50^. If no MDL Molfile could be obtained in this way, we instead downloaded the International Chemical Idenitifier (InChI) string via the mapping service of MetaCyc^54^, if a ChEBI ID was available. We then used the package Chem from RDKit^52^ with the MDL Molfiles or InChI strings as the input to calculate the 1024-dimensional binary ECFPs^30^ with a radius (number of iterations) of 3.

### Calculating task-specific fingerprints for the small molecules using graph neural networks

In addition to the pre-defined ECFPs, we also used a graph neural network (GNN) to calculate task-specific numerical representations for the small molecules. GNNs are neural networks that can take graphs as their input^55,56,57^. A molecule can be represented as a graph by interpreting the atoms and bonds of the molecule as nodes and edges, respectively. To provide the GNN with information about the small molecules, we calculated feature vectors for every bond and every atom in all molecules^27^. These features include bond type, atom type, atom mass, valence, atomic number, and atom charge. To input this information into a GNN, the graphs and the feature vectors are encoded with tensors and matrices. While a graph is processed by a GNN, all atom feature vectors are iteratively updated for a pre-defined number of steps by using information of neighboring bond and atom feature vectors. Afterwards, all atom feature vectors are pooled together by applying the element-wise mean to obtain a single graph representation. The dimension *D* of the updated atom feature vectors and of the final graph representation can be freely chosen; we chose *D* = 100. We trained and implemented a variant of GNNs called Directed Message Passing Neural Network (D-MPNN)^32^, using the Python package PyTorch^47^.

This small molecule representation was then concatenated with a small representation of an enzyme. To compute this enzyme representation, we performed principal component analysis (PCA)^58^ on the *ESM-1b* vectors (see below), selecting the first 50 principal components. The concatenated enzyme-small molecule vector was used as the input for a fully connected neural network (FCNN) with two hidden layers of size 100 and 32, which was trained for predicting whether the small molecule is a substrate for the enzyme. We trained the whole model (the GNN including the FCNN) end-to-end. Thereby, the model was forced to store task-specific and meaningful information in the graph representations. After training, we extracted a graph representation for every small molecule in our training set. For more details regarding training and implementation see our GitHub repository.

### Calculating enzyme representations

We used the *ESM-1b* model^2^ to calculate 1280-dimensional numerical representations of the enzymes. The *ESM-1b* model is a transformer network^37^ that takes amino acid sequences as its input and produces numerical representations of the sequences. First, every amino acid in a sequence is converted into a 1280-dimensional representation, which encodes the type of the amino acid and its position in the sequence. Afterwards, every representation is updated iteratively for 33 update steps by using information about the representation itself as well as about all other representations of the sequence using the attention mechanism^59^. The attention mechanism allows the model to selectively focus only on relevant amino acid representations to make updates^59^. During training ∼ 15% of the amino acids in a sequence are randomly masked and the model is trained to predict the type of the masked amino acids. The *ESM-1b* model was trained with ∼ 27 million proteins from the UniRef50 dataset^38^. To create a single representation for the whole enzyme, the element-wise mean of all updated amino acids representations in a sequence is calculated^2^. We created these representations for all enzymes in our dataset using the code and the trained *ESM-1b* model provided by the Facebook AI Research team on GitHub.

### Modifying the *ESM-1b* model to create task-specific enzyme representations

To create task-specific enzyme representations for our task of predicting enzyme-substrate pairs, we modified the *ESM-1b* model. For every input sequence, we added a representation for the whole enzyme in addition to the representations for all individual amino acids in the sequence. This representation is updated in the same way as the amino acid representations. The parameters of this modified model are initialized with the parameters of the trained *ESM-1b* model. After the last update layer of the model, i.e., after 33 update steps, we take the 1280-dimensional representation of the whole enzyme and concatenate it with a representation for a metabolite, the 1024-dimensional ECFP vector (see above). This concatenated vector is then used as the input for a fully-connected neural network (FCNN) with two hidden layers of size 256 and 32. The whole model was trained end-to-end for the binary classification task of predicting whether the added metabolite is a substrate for the given enzyme. This training procedure forced the model to store all necessary enzyme information for the prediction task in the enzyme representation. After training the modified model, we extracted the updated and task-specific representations, the *ESM-1b*_*ts*_ vectors, for all enzymes in our dataset.

We implemented and trained this model using the Python package PyTorch^47^. We trained the model with 287386 data points with phylogenetically inferred or experimental evidence for 2 epochs on 6 NVIDA DGX A100s, each with 40GB RAM. Training the model for more epochs did not lead to improved results. Because of the immense computational power and long training times, it was not possible to perform a systematic hyperparameter optimization. We chose hyperparameters after trying a few selected hyperparameter settings with values similar to the ones that were used for training the original *ESM-1b* model.

### Hyperparameter optimization of the gradient boosting models

To find the best hyperparameters for the gradient boosting models, we performed 5-fold cross-validations (CVs). To ensure a high diversity between all folds, we created the five folds in such a way that the same enzyme would not occur in two different folds. We used the Python package hyperopt^60^ to perform a random grid search for the following hyperparameters: learning rate, maximum tree depth, lambda and alpha coefficients for regularization, maximum delta step, minimum child weight, number of training epochs, and weight for negative data points. The last hyperparameter was added because our dataset is imbalanced; it allows the model to assign a lower weight to the negative data points during training. To ensure that our model is indeed not assigning too many samples to the over-represented negative class, we used a custom loss function that contains the False Negative Rate, *FNR*, and the False Positive Rate, *FPR*. Our loss function, 2 × *FNR*^2^ + *FPR*^1.3^, penalizes data points that are mistakenly assigned to the negative class stronger than data points that are mistakenly assigned to the positive class. After hyperparameter optimization, we chose the set of hyperparameters with the lowest mean loss during CV. We used the python package xgboost^40^ for training the gradient boosting models.

### Validating our model on two additional test sets

We compared the performance of ESP with two published models for predicting the substrate scope of single enzyme families. One of these models is a machine learning model for predicting the substrates of 12 different bacterial nitrilases^3^. Their dataset consists of 240 data points, where each of the 12 nitriliases was tested with the same 20 small molecules. This dataset was randomly split by Mou et al.^3^ into 80% training data and 20 % test data. We added all training data to our training set. After re-training, we validated our model performance on the test set from Ref.^3^.

The second model that we used for model comparison is a decision tree-based model, called GT-predict, for predicting the substrate scope of glycosyltransferases of plants^4^. As a training set, Yang et al.^4^ used 2847 data points with 59 different small molecules and 53 different enzymes, all coming from *Arabidopsis thaliana*. They used two independent test sets to validate model performance: one dataset with 266 data points with 7 enzymes from *Avena strigose* and 38 different small molecules, and a second dataset with 380 data points with 10 enzymes from *Lycium barbarum* and 38 different small molecules. We added all training data to our training set. After re-training, we validated ESP model performance on both test sets from Ref.^4^.

### Analyzing the effect of training set size

To analyze the effect of different training set sizes, we created eight different subsets of our training set with sizes ranging from 30% to 100% of the original training set size. To create these subsets, we first generated an enzyme list containing all enzymes of the training set in random order. To create the subsets, we extracted all training data points with enzymes that occur in the first 30%, 40%, …, 100% of the generated enzyme list. Afterwards, we re-trained our model on all different subsets of the training set and validated each version on our full test set.

### Statistical tests for model comparison

We tested if the difference in model performance between the two models with *ESM-1b* and *ESM-1b*_*ts*_ vectors is statistically significant. For this purpose, we used McNemar’s test^61^ (implemented in the Python package Statsmodels^62^), testing the null hypothesis that both models have a similar proportion of errors on our test set. We could reject the null hypothesis (*p <* 10^−28^), concluding that using *ESM-1b*_*ts*_ vectors leads to a statistically significant improvement.

We also tested if the differences in model performance between the three different splits of our test set with different enzyme sequence identity levels (0-40%, 40-60%, and 60-80%) are statistically significant. We used the non-parametric two-sided Mann–Whitney *U* test^63^ (implemented in the Python package SciPy^64^) to test the null hypothesis that the prediction errors for the different splits are equally distributed.

## Acknowledgements

We thank Veronica Maurino for insightful discussions. Computational support and infrastructure was provided by the “Centre for Information and Media Technology” (ZIM) at the University of Düsseldorf (Germany). We acknowledge financial support to MJL by the German Research Foundation (DFG) through CRC 1310, and, under Germany’s Excellence Strategy, through EXC2048/1 (Project ID:390686111), as well as through a grant by the Volkswagen Foundation under the “Life” initiative.

## Author contributions

SR implemented and trained the task-specific *ESM-1b*_*ts*_ model. AK designed the dataset and models and performed all other analyses. MKME conceived of the study. MJL supervised the study and acquired funding. AK, MKME, and MJL wrote the manuscript.

## References

[1] “UniProt: the universal protein knowledgebase in 2021”. In: Nucleic Acids Research 49.D1 (2021), pp. D480–D489.

[2] Alexander Rives et al. “Biological structure and function emerge from scaling unsupervised learning to 250 million protein sequences”. In: Proceedings of the National Academy of Sciences 118.15 (2021).

[3] Zhongyu Mou et al. “Machine learning-based prediction of enzyme substrate scope: Application to bacterial nitrilases”. In: Proteins: Structure, Function, and Bioinformatics 89.3 (2021), pp. 336–347.

[4] Min Yang et al. “Functional and informatics analysis enables glycosyltransferase activity prediction”. In: Nature chemical biology 14.12 (2018), pp. 1109–1117.

[5] Geoffrey M Cooper, Robert E Hausman, and Robert E Hausman. The cell: a molecular approach. Vol. 4. ASM press Washington, DC, 2007.

[6] Shelley D Copley. “Shining a light on enzyme promiscuity”. In: Current opinion in structural biology 47 (2017), pp. 167–175.

[7] Olga Khersonsky Tawfik and Dan S. “Enzyme promiscuity: a mechanistic and evolutionary perspective”. In: Annual review of biochemistry 79 (2010), pp. 471–505.

[8] Irene Nobeli, Angelo D Favia, and Janet M Thornton. “Protein promiscuity and its implications for biotechnology”. In: Nature biotechnology 27.2 (2009), pp. 157–167.

[9] Jose L Adrio and Arnold L Demain. “Microbial enzymes: tools for biotechnological processes”. In: Biomolecules 4.1 (2014), pp. 117–139.

[10] Songwei Wang et al. “Engineering a Synthetic Pathway for Gentisate in Pseudomonas Chlororaphis P3”. In: Frontiers in bioengineering and biotechnology 8 (2021), p. 1588.

[11] Ming-Cheng Wu et al. “Bioengineering natural product biosynthetic pathways for therapeutic applications”. In: Current opinion in biotechnology 23.6 (2012), pp. 931–940.

[12] Elzbieta Rembeza et al. “Discovery of Two Novel Oxidases Using a High-Throughput Activity Screen”. In: ChemBioChem 23.2 (2022), e202100510.

[13] Chelsea K Longwell, Louai Labanieh, and Jennifer R Cochran. “High-throughput screening technologies for enzyme engineering”. In: Current opinion in biotechnology 48 (2017), pp. 196–202.

[14] Gary W Black et al. “A high-throughput screening method for determining the substrate scope of nitrilases”. In: Chemical Communications 51.13 (2015), pp. 2660–2662.

[15] Nicki Skafte Detlefsen, Søren Hauberg, and Wouter Boomsma. “Learning meaningful representations of protein sequences”. In: Nature Communications 13.1 (Apr. 2022), p. 1914. ISSN: 2041-1723. DOI: 10.1038/s41467-022-29443-w.

[16] Dante A Pertusi et al. “Predicting novel substrates for enzymes with minimal experimental effort with active learning”. In: Metabolic engineering 44 (2017), pp. 171–181.

[17] Marc Röttig, Christian Rausch, and Oliver Kohlbacher. “Combining structure and sequence information allows automated prediction of substrate specificities within enzyme families”. In: PLoS computational biology 6.1 (2010), e1000636.

[18] Marc G Chevrette et al. “SANDPUMA: ensemble predictions of nonribosomal peptide chemistry reveal biosynthetic diversity across Actinobacteria”. In: Bioinformatics 33.20 (2017), pp. 3202–3210.

[19] Jae Yong Ryu, Hyun Uk Kim, and Sang Yup Lee. “Deep learning enables high-quality and high-throughput prediction of enzyme commission numbers”. In: Proceedings of the National Academy of Sciences 116.28 (2019), pp. 13996–14001. DOI: 10.1073/pnas.1821905116. eprint: https://www.pnas.org/doi/pdf/10.1073/pnas.1821905116.

[20] Yu Li et al. “DEEPre: sequence-based enzyme EC number prediction by deep learning”. In: Bioinformatics 34.5 (Oct. 2017), pp. 760–769. ISSN: 1367-4803. DOI: 10.1093/bioinformatics/btx680. eprint: https://academic.oup.com/bioinformatics/article-pdf/34/5/760/25117683/btx680.pdf.

[21] Theo Sanderson et al. “ProteInfer: deep networks for protein functional inference”. In: bioRxiv (2021). DOI: 10.1101/2021.09.20.461077. eprint: https://www.biorxiv.org/content/early/2021/10/06/2021.09.20.461077.full.pdf.

[22] Maxwell L. Bileschi et al. “Using deep learning to annotate the protein universe”. In: Nature Biotechnology (Feb. 2022). ISSN: 1546-1696. DOI: 10.1038/s41587-021-01179-w.

[23] Elzbieta Rembeza and Martin KM Engqvist. “Experimental investigation of enzyme functional annotations reveals extensive annotation error”. In: bioRxiv (2020). DOI: 10.1101/2020.12.18.423474. eprint: https://www.biorxiv.org/content/early/2020/12/19/2020.12.18.423474.full.pdf.

[24] Hakime Öztürk, Arzucan Özgür, and Elif Ozkirimli. “DeepDTA: deep drug–target binding affinity prediction”. In: Bioinformatics 34.17 (2018), pp. i821–i829.

[25] Qingyuan Feng et al. “Padme: A deep learning-based framework for drug-target interaction prediction”. In: arXiv preprint 1807.09741 (2018).

[26] Mostafa Karimi et al. “DeepAffinity: interpretable deep learning of compound–protein affinity through unified recurrent and convolutional neural networks”. In: Bioinformatics 35.18 (2019), pp. 3329–3338.

[27] Alexander Kroll et al. “Deep learning allows genome-scale prediction of Michaelis constants from structural features”. In: PLoS biology 19.10 (2021), e3001402.

[28] Feiran Li et al. “Deep learning based kcat prediction enables improved enzyme constrained model reconstruction”. In: bioRxiv (2021). DOI: 10.1101/2021.08.06.455417. eprint: https://www.biorxiv.org/content/early/2021/08/08/2021.08.06.455417.full.pdf.

[29] David Weininger. “SMILES, a chemical language and information system. 1. Introduction to methodology and encoding rules”. In: Journal of Chemical Information and Computer Sciences 28.1 (1988), pp. 31–36.

[30] David Rogers and Mathew Hahn. “Extended-connectivity fingerprints”. In: Journal of Chemical Information and Modeling 50.5 (2010), pp. 742–754.

[31] Jie Zhou et al. “Graph neural networks: A review of methods and applications”. In: AI Open 1 (2020), pp. 57–81.

[32] Kevin Yang et al. “Analyzing learned molecular representations for property prediction”. In: Journal of chemical information and modeling 59.8 (2019), pp. 3370–3388.

[33] Ethan C Alley et al. “Unified rational protein engineering with sequence-based deep representation learning”. In: Nature methods 16.12 (2019), pp. 1315–1322.

[34] Yuting Xu et al. “Deep dive into machine learning models for protein engineering”. In: Journal of chemical information and modeling 60.6 (2020), pp. 2773–2790.

[35] Jacob Devlin et al. “Bert: Pre-training of deep bidirectional transformers for language understanding”. In: arXiv preprint 1810.04805 (2018).

[36] Emily C Dimmer et al. “The UniProt-GO annotation database in 2011”. In: Nucleic acids research 40.D1 (2012), pp. D565–D570.

[37] Ashish Vaswani et al. “Attention is all you need”. In: Advances in neural information processing systems. 2017, pp. 5998–6008.

[38] Baris E Suzek et al. “UniRef clusters: a comprehensive and scalable alternative for improving sequence similarity searches”. In: Bioinformatics 31.6 (2015), pp. 926–932.

[39] Ahmed Elnaggar et al. “ProtTrans: Towards Cracking the Language of Lifes Code Through Self-Supervised Deep Learning and High Performance Computing”. In: IEEE transactions on pattern analysis and machine intelligence PP (July 2021). ISSN: 0162-8828. DOI: 10.1109/tpami.2021.3095381.

[40] Tianqi Chen and Carlos Guestrin. “Xgboost: A scalable tree boosting system”. In: Proceedings of the 22nd acm sigkdd international conference on knowledge discovery and data mining. 2016, pp. 785–794.

[41] Davide Chicco and Giuseppe Jurman. “The advantages of the Matthews correlation coefficient (MCC) over F1 score and accuracy in binary classification evaluation”. In: BMC genomics 21.1 (2020), pp. 1–13.

[42] Weidong Tian and Jeffrey Skolnick. “How well is enzyme function conserved as a function of pairwise sequence identity?” In: Journal of molecular biology 333.4 (2003), pp. 863–882.

[43] Antje Chang et al. “BRENDA, the ELIXIR core data resource in 2021: new developments and updates”. In: Nucleic Acids Research 49.D1 (Jan. 2021), pp. D498–D508. ISSN: 0305-1048. DOI: 10.1093/nar/gkaa1025.

[44] John Jumper et al. “Highly accurate protein structure prediction with AlphaFold”. In: Nature 596.7873 (2021), pp. 583–589.

[45] Minkyung Baek et al. “Accurate prediction of protein structures and interactions using a three-track neural network”. In: Science 373.6557 (2021), pp. 871–876. DOI: 10.1126/science.abj8754. eprint: https://www.science.org/doi/pdf/10.1126/science.abj8754.

[46] Guido Van Rossum and Fred L. Drake. Python 3 Reference Manual. Scotts Valley, CA: CreateSpace, 2009. ISBN: 1441412697.

[47] Adam Paszke et al. “Pytorch: An imperative style, high-performance deep learning library”. In: Advances in neural information processing systems 32 (2019), pp. 8026–8037.

[48] “The Gene Ontology resource: enriching a GOld mine”. In: Nucleic Acids Research 49.D1 (2021), pp. D325–D334.

[49] Parit Bansal et al. “Rhea, the reaction knowledgebase in 2022”. In: Nucleic acids research (2021).

[50] Minoru Kanehisa and Susumu Goto. “KEGG: kyoto encyclopedia of genes and genomes”. In: Nucleic acids research 28.1 (2000), pp. 27–30.

[51] Sunghwan Kim et al. “PubChem in 2021: new data content and improved web interfaces”. In: Nucleic acids research 49.D1 (2021), pp. D1388–D1395.

[52] Greg Landrum et al. RDKit: Open-source cheminformatics. http://www.rdkit.org. 2006.

[53] Limin Fu et al. “CD-HIT: accelerated for clustering the next-generation sequencing data”. In: Bioinformatics 28.23 (2012), pp. 3150–3152.

[54] Ron Caspi et al. “The MetaCyc database of metabolic pathways and enzymes-a 2019 update”. In: Nucleic acids research 48.D1 (2020), pp. D445–D453.

[55] Steven Kearnes et al. “Molecular graph convolutions: moving beyond fingerprints”. In: Journal of Computer-Aided Molecular Design 30.8 (2016), pp. 595–608.

[56] David K Duvenaud et al. “Convolutional networks on graphs for learning molecular fingerprints”. In: Advances in Neural Information Processing Systems. 2015, pp. 2224–2232.

[57] Jie Zhou et al. “Graph neural networks: A review of methods and applications”. In: arXiv preprint (2018), 1812.08434.

[58] Ian Jolliffe. “Principal component analysis”. In: Encyclopedia of statistics in behavioral science (2005).

[59] Dzmitry Bahdanau, Kyunghyun Cho, and Yoshua Bengio. “Neural machine translation by jointly learning to align and translate”. In: arXiv preprint 1409.0473 (2014).

[60] James Bergstra, Daniel Yamins, and David Cox. “Making a science of model search: Hyperparameter optimization in hundreds of dimensions for vision architectures”. In: International conference on machine learning. PMLR. 2013, pp. 115–123.

[61] Thomas G Dietterich. “Approximate statistical tests for comparing supervised classification learning algorithms”. In: Neural computation 10.7 (1998), pp. 1895–1923.

[62] Skipper Seabold and Josef Perktold. “Statsmodels: Econometric and statistical modeling with python”. In: Proceedings of the 9th Python in Science Conference. Vol. 57. Austin, TX. 2010, p. 61.

[63] Patrick E McKnight and Julius Najab. “Mann-Whitney U Test”. In: The Corsini encyclopedia of psychology (2010), pp. 1–1.

[64] Pauli Virtanen et al. “SciPy 1.0: fundamental algorithms for scientific computing in Python”. In: Nature methods 17.3 (2020), pp. 261–272.

