## Supplementary Tables & Figures for "The substrate scopes of enzymes: a general prediction model based on machine and deep learning"

### Supplementary Information

#### Supplementary Tables S1-S2

**Table S1.** Results of hyperparameter optimizations of the gradient boosting models for all four combinations of small molecule representations (ECFPs and GNN generated fingerprints) and enzyme representations (*ESM-1b* and *ESM-1b*$_{ts}$ vectors). The hyperparameter optimizations were performed with 5-fold cross-validation on the training set.

| | **mean ROC-AUC (CV)** | **learning rate** | **max. delta step** | **max. depth** | **min. child weight** | **num. of trees** | **alpha coeff.** | **beta coeff.** | **weight** |
|---|---|---|---|---|---|---|---|---|---|
| ***ESM-1b* & ECFP** | 0.937 | 0.127 | 3.08 | 13 | 2.69 | 333 | 1.43 | 0.12 | 0.114 |
| ***ESM-1b*$_{ts}$ & ECFP** | 0.950 | 0.316 | 1.77 | 10 | 1.38 | 343 | 0.53 | 3.74 | 0.262 |
| ***ESM-1b* & GNN** | 0.946 | 0.187 | 3.75 | 10 | 0.40 | 367 | 0.89 | 4.89 | 0.142 |
| ***ESM-1b*$_{ts}$ & GNN** | 0.954 | 0.184 | 3.27 | 13 | 3.19 | 314 | 0.48 | 2.62 | 0.126 |

**Table S2.** Results of validating the ESP model on the test sets from Yang et al.[4] and Mou et al.[3] without adding any new training data to our training set.

| | **ROC-AUC score** | **Accuracy** | **MCC** |
|---|---|---|---|
| ***Yang et al. Avena strigosa*** | 0.59 | 68% | 0.12 |
| ***Yang et al. Lycium barbarum*** | 0.56 | 66% | 0.01 |
| ***Mou et al.*** | 0.41 | 0.5% | 0.00 |

#### Supplementary Figures S1-S2

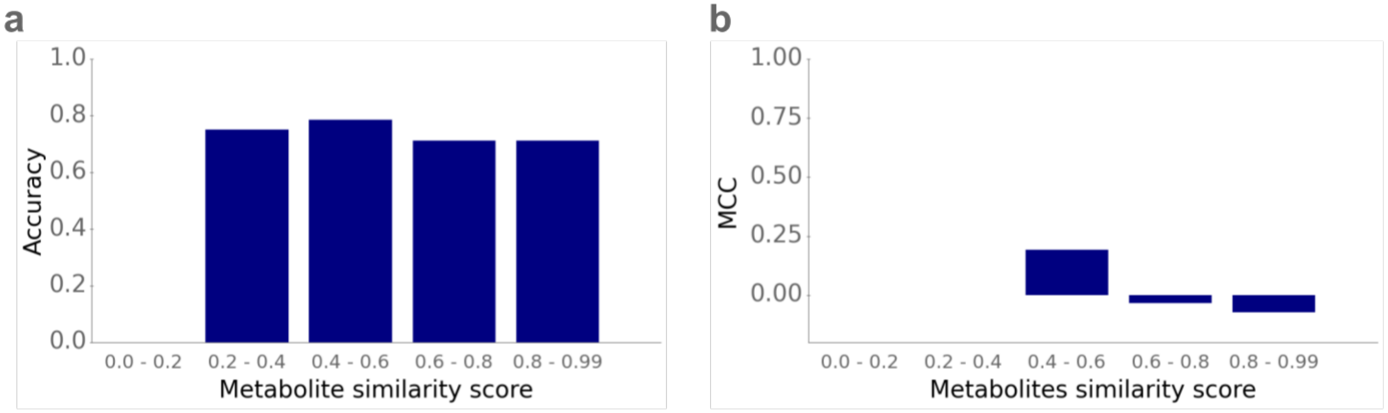


**Figure S1. Effect of the metabolite similarity score on model performance.** For all small molecules in the test set that do not also occur in the training set, we calculated the maximal pairwise similarity score across all small molecules in the training set. The similarity score is a value between 0 and 1, where a higher value indicates higher similarity between a pair of metabolites. We divided all test data points with small molecules that do not occur in the training set into five subsets dependent on their maximal similarity scores. **(a)** shows the accuracy and **(b)** shows the MCC for each subset.

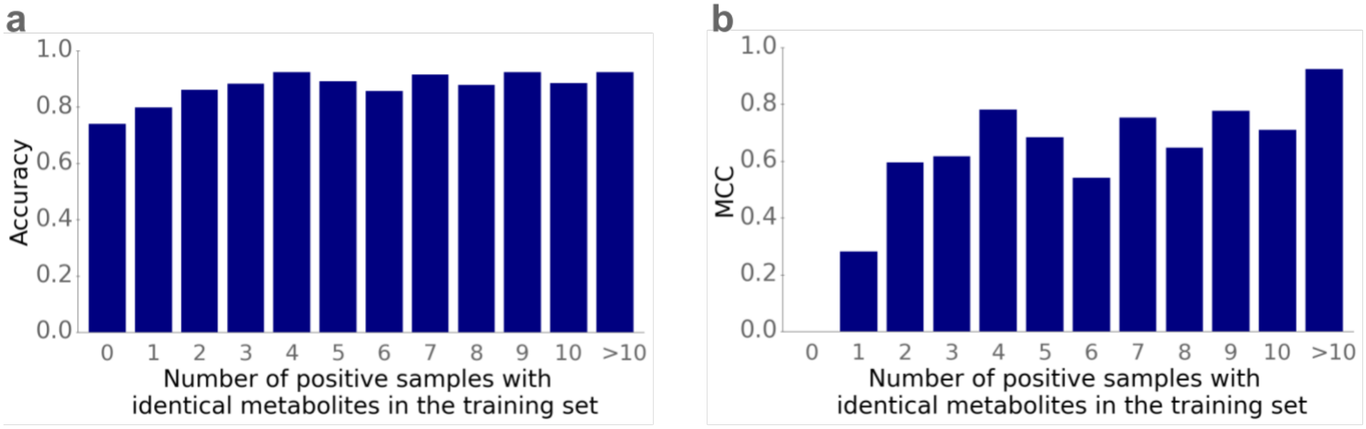


**Figure S2. Effect of the number of identical substrates in the training set on model performance.** We grouped small molecules by how often they occur as substrates among all positive data points in the training set. **(a)** shows the accuracy and **(b)** shows the MCC for each group.
